# Does What You Eat Affect How You Mate? Disentangling the Interactions Between Diet-Induced Phenotypic Plasticity and Adult Reproductive Strategies in Black Soldier Flies

**DOI:** 10.1101/2024.03.03.583236

**Authors:** Qi-Hui Zhang, Keng Hee Ng, Wells Shijian Chin, Yong Jen Tang, Jielin Lin, Nalini Puniamoorthy

## Abstract

Phenotypic plasticity enables organisms to response to environmental variations by generating a range of phenotypes from a single genotype. In holometabolous insects, traits that influence larval plasticity may hold relevance for adult life history strategies. We present a comprehensive investigation into phenotypic plasticity in black soldier flies, a species known for its efficient waste-to-biomass conversion in the larval stage. Here, we document adult sex-specific plastic responses and reproductive strategies shaped by larval diets. We examined traits including adult body size, reproductive organ development, sperm length, mating behaviours, egg production and other life history parameters across different treatments. Our findings reveal notable sex-specific differences in phenotypic plasticity, with females showing increased plasticity in reproductive investment. Furthermore, males and females differed starkly in allometric growth and weight ratio of reproductive organs. Diets that facilitated longer male lifespans also prompted earlier male emergence suggesting an interplay between lifespan and degree of protandry. This maximizes the overlap of male and female lifespans, thereby enhancing mating success in diverse environmental conditions. Our results reveal plastic responses in mating behaviours, where diets producing smaller adults, smaller reproductive organs, and shorter sperm correlated with significantly enhanced mating effort and performance. This study highlights the complex interactions between nutrition, development, and reproductive strategies, and has significant implications for the insect bioconversion industries.

## Introduction

Phenotypic plasticity, defined as the capacity to generate a range of phenotypes from a single genotype in response to environmental variations, holds relevance for many insect species (Derek A. Roff 2009). One of the most pivotal environmental variables that induces phenotypic plasticity in holometabolous insects is larval diet quality (Baleba et al. 2019, Kröncke and Benning 2023). Early larval stages are especially susceptible to environmental cues such as nutritional variation (Zhang et al. 2023a), which can in turn shape their adult developmental outcomes (Rehman and Varghese 2021). In addition, the influence of these early-life dietary factors can be enduring, having lasting effects on adult growth, behaviour, and physiological functions, even when the initial environmental conditions are no longer present (Buchanan et al. 2022, Liu et al. 2022). For instance, nutritional intake during the larval stage can induce plasticity in the horn length of adult male dung beetles (Reaney and Knell 2015), a trait that is associated with reproductive success via intrasexual selection (Emlen et al. 2005). Moreover, females and males could respond differently to environmental variation. This sex difference in phenotypic plasticity can provide insights into the proximate mechanisms of the developmental processes mediating adult sexual size dimorphism (SSD) (Stillwell et al. 2009).

Black Soldier Flies (BSF, *H. illucens* L., Diptera: Stratiomyidae) present an excellent model for examining the impact of larval diet on adult phenotypic plasticity. Known for their remarkable versatility in decomposing organic wastes, BSF larvae can thrive on a diverse array of organic substrates, from agricultural and restaurant wastes to byproducts of food processing industries, and even to wastewater, municipal and animal manure (Purkayastha and Sarkar 2022). Extensive research has already been conducted on the plastic responses during the larval stage (Scala et al. 2020, Zhang et al. 2023b), primarily because it is directly linked to the yields and quality of end products like animal feed and biodiesel (Wang et al. 2017, Lu et al. 2022, Siva Raman et al. 2022). However, a crucial yet underexplored dimension of BSF biology is how the nutritional diversity of larval diets affects adult phenotypes, particularly with respect to traits associated with reproduction (Lemke et al. 2023, Meneguz et al. 2023).

Adult BSFs do not require additional feeding post-metamorphosis, relying instead on nutrient reserves accumulated during their larval stage (Amrul et al. 2022). This fact magnifies the potential importance of larval diet in shaping adult phenotypes, especially concerning the phenotypic plasticity in reproductive traits. In addition, plasticity can be adaptive maladaptive or neutral, depending on its potential effects on the final fitness of individuals (Crowther et al. 2023). This perspective underscores the need for a more holistic approach to studying BSF plasticity, one that considers the adaptiveness of plastic responses across a spectrum of reproductive traits to better understand their effects of reproductive strategies.

Previous studies on adult BSF plasticity have largely focused on specific parameters such as oviposition and total mating pairs (Tomberlin and Sheppard 2002, Heussler et al. 2018), mating success and egg hatching rate (Liu et al. 2022), longevity (Thinn and Kainoh 2022, Harjoko et al. 2023), and life-history traits and sex ratios (Gobbi et al. 2013). These studies often vary in the strain of BSF used, the rearing environments and experimental designs. There still exists a gap in the literature that comprehensively investigates the full spectrum of reproductive traits in both male and female BSFs.

Our study aims to address this research gap by providing a nuanced exploration of phenotypic plasticity in adult BSFs, particularly focusing on the impact of larval diet on an extensive range of reproductive traits across both sexes. While larval stage traits such as larval growth and development were recorded, our primary focus is on adult traits that could influence reproductive success. These include variations in protandry (early emergence), adult longevity, female egg production, mating behaviours, adult body size, and reproductive organ development—gonadal development (ovaries/testes) and accessory glands in both sexes, as well as sperm length in males. Moreover, our research also aims to compare the phenotypic plasticity of male and female adults in terms of plastic degree and pattern, thereby shedding light on sex-specific variations in phenotypic plasticity. Through this approach, we seek to deepen our understanding of how larval developmental environments sculpt adult phenotypes and reproductive strategies. Such findings hold significant implications for the long-term sustainability of BSF colonies, and by extension, the viability of the BSF industry.

## Materials and Methods

### Experimental insects

A lab-adapted genetic line of BSF was used in this study. This strain was established in 2018 through crossbreeding between Southeast Asian and North American populations (Eugene 2019) and has been maintained on food waste from residential dining halls (labelled as HF in this study) at the National University of Singapore (NUS) BSF facility (1°17’45.2“N, 103°46’44.5”E) (for details see Zhang et al. 2023b). For this experiment, the eggs from line E were collected and transferred to the nursery for hatching. The newly hatched larvae were reared on chicken feed for 5 days before being transferred to the experimental substrates. All experiments were conducted at room temperature ∼28.3 °C with an environmental humidity of ∼85.3% relative humidity (%RH).

### Larval meals

Four experimental larval meals were utilized in this study alongside commercial chicken feed (CF, PK Agro-Industrial Products M Sdn. Bhd., Johor, Malaysia) as the control meal (Bosch et al. 2020) (Table S1-1). The meals were categorized based on their origin: VF (Vegetable and Fruit Waste), AB (Agricultural Byproduct), HF (Campus Hall Food Waste), and RN (Rice and Noodles), representing diverse components of urban food waste streams. The food waste materials were obtained from two different sources: (1) Tiong Lam Supplies Pte. Ltd, Singapore, a Singaporean food waste management company; (2) residential dining halls of the NUS. The nutrient composition of the meals was analysed, including crude protein, crude fat, crude carbohydrate, and ash content (Table S1). Furthermore, the amino acid profiles of the larval meals, featuring Asp, Thr, Ser, Glu, Gly, Ala, Cystine, Val, Met, Ile, Leu, Tyr, Phe, Lys, His, Arg, and Pro, are detailed in Table S1-2.

### Larval rearing setup

Five-day old larvae were weighed using high-precision analytical balance (0.000001g, QUINTIX35-1S, Sartorius, Germany) and then counted under microscope (Olympus SZX2-FO, Spectra, USA) to estimate the individual weight (10 replicates). For each diet treatment, 250 neonates were weighed and placed in a plastic container (10 x 12 x 5cm) containing a 350 g meal with 70%-80% water content. During the rearing phase, these containers, equipped with netted lids (100 Mesh), were placed within a larger net cage. Four biological replicates were maintained for each treatment setup.

### Larval development, adult emergence, and longevity

The number of prepupae were recording from day 12 to day 19 of larval rearing. The individual prepupal weight was calculated by dividing the cumulative weight with the total number of prepupae. Pupae were individually placed in cup containers to ensure the collection of virgin flies. These containers were inspected daily; newly emerged adults were promptly identified and sexed according to their terminal abdominal morphology (Julita et al. 2020). Each container was marked with the emergence date and the sex of the adult fly to determine the degree of protandry.

To evaluate the longevity of male and female BSFs, individuals that emerged on the same day were randomly allocated and separated by sex into nylon cages (20-30 flies per cage) at four days post-emergence. Cages were equipped with a damp sponge to supply moisture. Daily inspections were conducted to record and remove any dead flies.

### Mating and mating behaviours

For each dietary treatment group, 20 virgin males and 20 females aged between three to four days were selected randomly and introduced into mating enclosures. To induce mating behaviour, artificial illumination approximating 7950 Lux was provided. Following a 5-second acclimation in darkness, achieved by enclosure with a black bag, a 15-minute session was video recorded to capture the mating behaviours.

Subsequent video analysis was employed to count the number of male mating attempts and the occurrence of successful copulation events. Mating attempts were quantified by tallying instances where a male mounted a female for a duration exceeding one second. A successful event was identified as successful upon observation of reciprocal copulation, characterized by the alignment and connection of genitalia in an antipodal configuration (Julita et al. 2020). Furthermore, female resistance was measured from the start of the male’s mounting to the point of genital coupling, providing an index of female receptivity.

### Adult morphological traits, reproductive organ development and sperm length

Random samples of ten virgin females and males were selected from each replicate at four days post-eclosion and preserved at -20L for subsequent analysis. Adult specimens were weighed with an analytical balance (QUINTIX35-1S, Sartorius, Germany, precision of 0.00001g). Morphological parameters, including head width and thoracic scutum length, were meticulously measured under a dissecting microscope (Olympus SZX10 FO, USA). Dissections were conducted in 1x PBS buffer (pHL6.5) under the microscope (Figure 2a).

Gonads, specifically male testes and accessory glands, as well as female ovaries and accessory glands, were dissected out and placed on aluminium foil. Any residual buffer was gently removed with a Kimwipe (KIMTECH) to ensure accurate subsequent weighing of the tissues.

During the dissection of each male BSF, the seminal vesicles were carefully pierced with forceps, allowing the sperm to be collected onto a microscope slide mixed with one drop of 1x PBS. The slide was then dried at 60L. Following this, all dried slides were fixed using a Methanol: Acetic acid solution (3:1 ratio) and subsequently stained with DAPI staining solution. The slides were left to dry in a dark environment before being stored in slide boxes for later analysis. Using the cellSens imaging software, sperm head and tail measurements were taken from ten sperms on each slide under a fluorescence microscope (Olympus SZX10 FO, USA) with a 20× objective.

### Statistical Analysis

Dataset distribution was assessed for normality using QQ-plots, followed by the application of Bartlett’s test to evaluate the homogeneity of variances across groups. Analysis of variance (one-way ANOVA) and Kruskal-Wallis test were used to determine the effects of rearing meals on prepupae development and multiple adult traits. Differences were regarded as significant at p-value < 0.05 for all tests. If significant differences were found, post-hoc tests such as Tukey-Kramer HSD Test and Nemenyi-Damico-Wolfe-Dunn test were performed to identify meal types with significantly different means and categorise them into groups.

To evaluate and compare the sex-specific plasticity in body size parameters and reproductive organ parameters of BSF adults, Reduced Major Axis (RMA) regression analysis was conducted using log-transformed data for male and female BSFs. The degree of sex-specific plasticity was quantified through two calculation methods: the coefficient of variation-total and the max-min medians index (Valladares et al. 2006). Additionally, with log-transformed data, a linear regression model was constructed to explore the allometric scaling relationships between body weight and gonad weight (ovaries and testes), and between body weight and accessory gland weight in female and male BSFs.

Larvae reared on the VF diet experienced a significantly high adult mortality rate, with the majority either failing to emerge fully or emerging with abnormal morphology and subsequently dying (Figure S1). Consequently, the VF group was omitted from the downstream analysis of adult traits.

All analyses were conducted using RStudio version 2023.06.0+421 (RStudio Team, 2023) and data were visualized with ggplot2 package in R.

## Results

### Diet-Induced Plasticity in Larval Growth and Development

Diet influenced BSF larval growth across substrates, with CF and HF diets leading to the heaviest prepupae, heavier than those on AB and significantly more than those on RN. VF-fed larvae were the lightest, nearly three times lighter than CF and HF groups (details in Table S2). Larval development rates also varied by diet; CF diet larvae quickly progressed to prepupae, particularly from Day 12 to 15. While AB and HF diets initially showed similar prepupal rates, HF’s prepupae numbers overtook AB after Day 15, peaking by Day 19. Conversely, VF and RN diets led to slower development, with VF and RN larvae transitioning to prepupae by Days 17 and 19, respectively (details in Table S2).

### Sexual Dimorphism in Phenotypic Plasticity

Figure 1a compares phenotypic plasticity between BSF females and males using RMA regression, focusing on body size and reproductive organ parameters, respectively. Notably, the RMA slope absolute values for body weight, head width, and scutum length are less than 1 (0.74, 0.84, and 0.78), indicating that in these traits, males display a higher degree of plasticity than females (Figure 1a). In contrast, the RMA slope values for GSI and the accessory gland to gonad ratio are significantly larger than 1 (107.11 and 234.81), revealing notably higher plasticity in female reproductive traits (Figure 1b). These findings are also supported by quantitative estimates of phenotypic plasticity, with body weight displaying higher plasticity degree in males, while reproductive parameters exhibiting converse trends of plasticity degree between the sexes (see Table 1).

**Figure 1.**
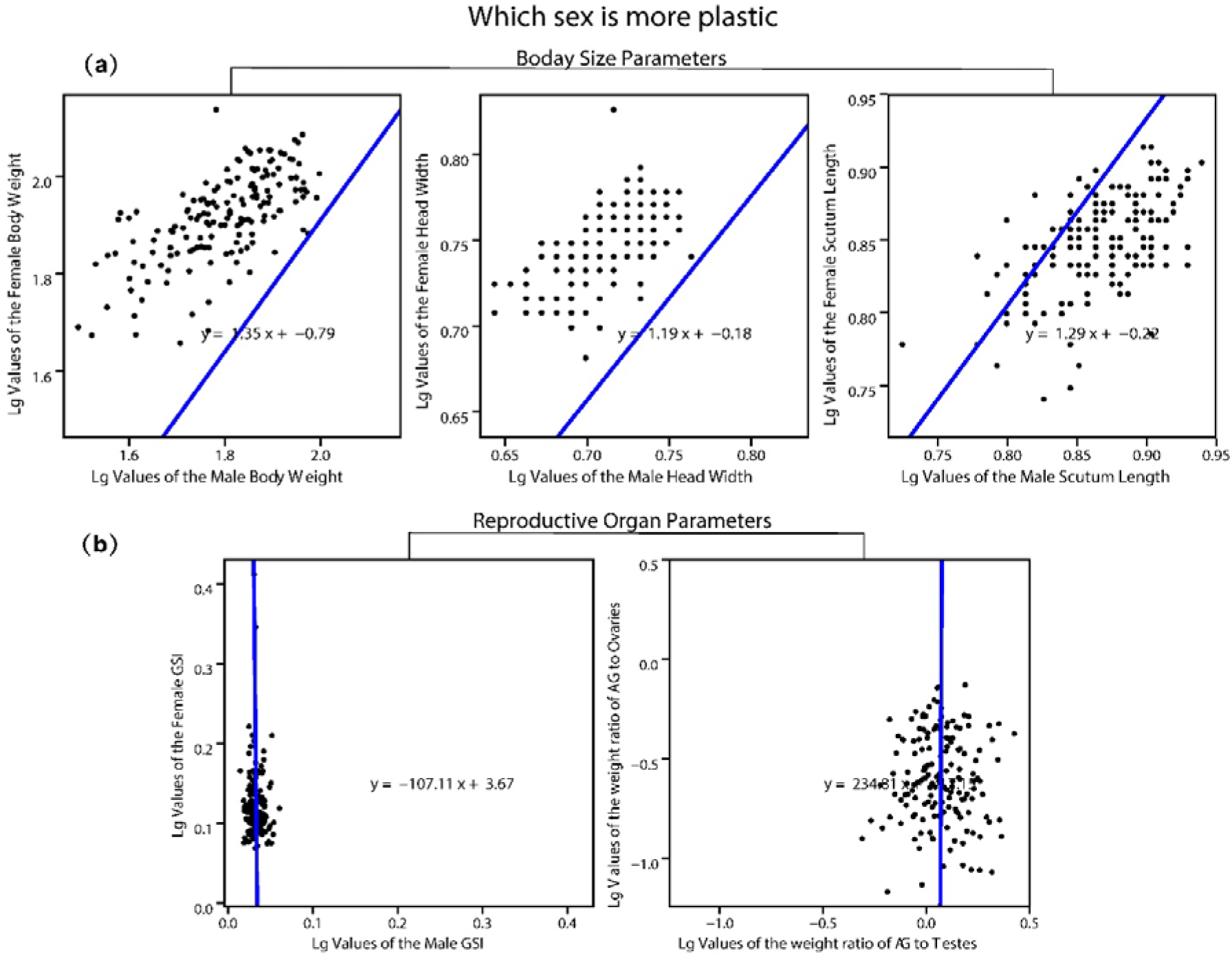
Plasticity Degree Comparison in Female and Male Body Size Parameters and Reproductive Organ Parameters. This figure presents the Reduced Major Axis (RMA) regression analysis, illustrating log-transformed values for: (a) body size parameters with male data on the x-axis and female data on the y-axis; and (b) Gonadosomatic Index (GSI) and the weight ratio of accessory glands to gonads, again with male data on the x-axis and female data on the y-axis. RMA slopes with absolute values greater than 1 indicate higher variation in female parameters compared to male, while slopes less than 1 suggest higher variation in males.

**Table 1.** Quantitative estimators of phenotypic plasticity and phenotypic inertia of the two BSF lines.

| Estimators of plasticity | Calculation<br>(based on lg values) | Results (absolute values) |  |
| --- | --- | --- | --- |
| Coefficient of variation-total | standard deviation/mean | adult body weight | $P_F (4.77) < P_M (6.21)$ |
| | | GSI | $P_F (35.20) > P_M (7.02)$ |
| | | AG to gonad ratio | $P_F (37.23) > P_M (7.02)$ |
| Max-min medians Index | (Maximum median-minimum median)/<br>maximum median | adult body weight | $P_F (7.95) < P_M (12.43)$ |
| | | GSI | $P_F (17.94) > P_M (3.32)$ |
| | | AG to gonad ratio | $P_F (77.01) > P_M (53.23)$ |

**Note:** $P_F$ and $P_M$ represent the quantified plasticity for females and males respectively. Three parameters phenotypic plasticity were quantified: adult body weight, gonadosomatic index (GSI), and the weight ratio of accessory glands to gonads in both females and males.

### Diet-Induced Plasticity in Body Size, Reproductive Organ Weight in BSF adults

In females, the impact of diet on body weight, head width, and scutum length was consistent. Females reared on HC and CF diets developed into the largest adults, followed by those from the AB group, while RN-fed larvae matured into the smallest females (Table 2). In males, larval diets influenced body weight and head width similarly to females. However, males reared on CF diet had longer scutum lengths compared to those reared on HF (Table 2). Larvae reared on the CF diet resulted in adult females with heavier accessory glands and ovaries (Figure 2a), whereas males from the HF diet matched the CF group in accessory gland and testes weights (Figure 2b). Across both sexes, the RN diet correlated with the lowest weights in accessory glands and reproductive organs. (Figure 2). In females, the ratio of accessory gland weight to ovary weight was significantly higher for those reared on the CF diet compared to the RN diet, which in turn had the lowest investment in accessory glands (Figure S2). Meanwhile, male BSFs showed no significant differences in the accessory gland to testes weight ratio when comparing across the various larval diet treatments (Figure S2).

**Table 2.** Comparison of Body Size Parameters in Male and Female BSF Across Various Diets.

| Meal | Female |  |  | Male |  |  |
| --- | --- | --- | --- | --- | --- | --- |
|  | Body weight<br>(mg) | Head width<br>(mm) | Scutum length<br>(mm) | Body weight<br>(mg) | Head width<br>(mm) | Scutum length<br>(mm) |
| AB | 82.15 ± 8.71 b | 5.56 ± 0.24 b | 7.08 ± 0.28 b | 60.70 ± 7.49 b | 5.10 ± 0.16 b | 7.24 ± 0.30 c |
| CF | 96.73 ± 12.61 a | 5.82 ± 0.17 a | 7.38 ± 0.45 a | 78.15 ± 10.41 a | 5.43 ± 0.13 a | 7.97 ± 0.30 a |
| HF | 93.26 ± 14.50 a | 5.76 ± 0.21 a | 7.31 ± 0.42 a | 74.32 ± 10.88 a | 5.37 ± 0.23 a | 7.72 ± 0.43 b |
| RN | 65.38 ± 10.89 c | 5.29 ± 0.18 c | 6.50 ± 0.43 c | 46.04 ± 8.17 c | 4.84 ± 0.20 c | 6.60 ± 0.41 d |

**Note:** The values represent mean ± standard error, and the letters behind the values represent significant difference according to TukeyHSD test and Kruskal test.

**Figure 2:**
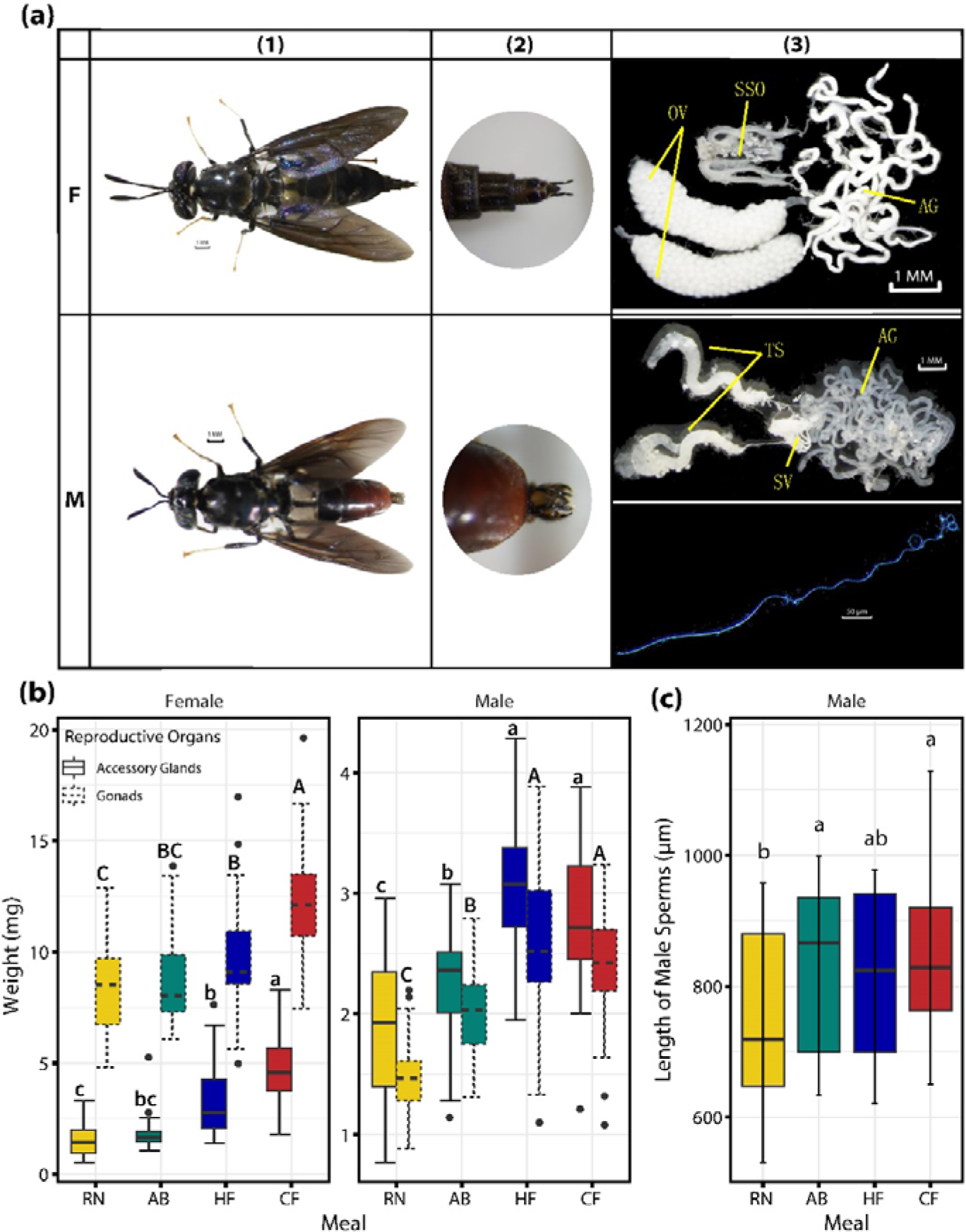
Variation in Reproductive Organs of Male and Female BSFs as well as Variation in Sperm Length. (a) representative images of adult BSFs, with ’F’ denoting females (top row) and ’M’ for males (bottom row). The first column (1) shows the whole body, the second column (2) displays the genitalia, and the third column (3) presents the reproductive organs (OV: ovary, AG: accessory glands, SSO: sperm storage organ, TS: testes, SV: seminal vesical) for both sexes and a male sperm; (b) the weight of the ovaries and accessory glands in female BSFs and the weight of the testes and accessory glands in male BSFs; and (c) the sperm length in male BSFs. Diets are classified by colour. In (b), different reproductive organs are represented by various box line types. Lowercase letters indicate significant differences in accessory gland weights and sperm length across diets, while uppercase letters mark significant differences in testes and ovaries weights (Kruskal-Wallis test, Bonferroni correction, P < 0.05).

### Diet-Induced Plasticity in Sperm Length and Allometric Scaling in BSF Males

In this study, a total of 1500 sperm samples were measured from 150 males reared on four different treatments to explore the impact of larval diet on sperm morphometrics. Our findings indicated that within each group, the total sperm length can vary drastically, from approximately 600 to over 1000 μm (Figure 2c), and head length can vary from about 15 to 21 μm (Figure S3). Males from the RN dietary group displayed notably shorter total sperm, as shown in Figure 3c. However, no significant difference in sperm length parameters were observed among males across all the dietary treatments. Interestingly, a negative correlation was observed between sperm length and body size parameters in RN males (Figure S4). Specifically, increases in scutum length and head width were inversely related to sperm length, with allometric coefficients of -1.2 and -0.37, respectively (Figure S4). Conversely, CF males demonstrated a significant positive allometric relationship between sperm length and scutum length (allometric coefficient =1.4). Across all groups, a hypoallometric pattern was noted with respect to head width and sperm length.

**Figure 3.**
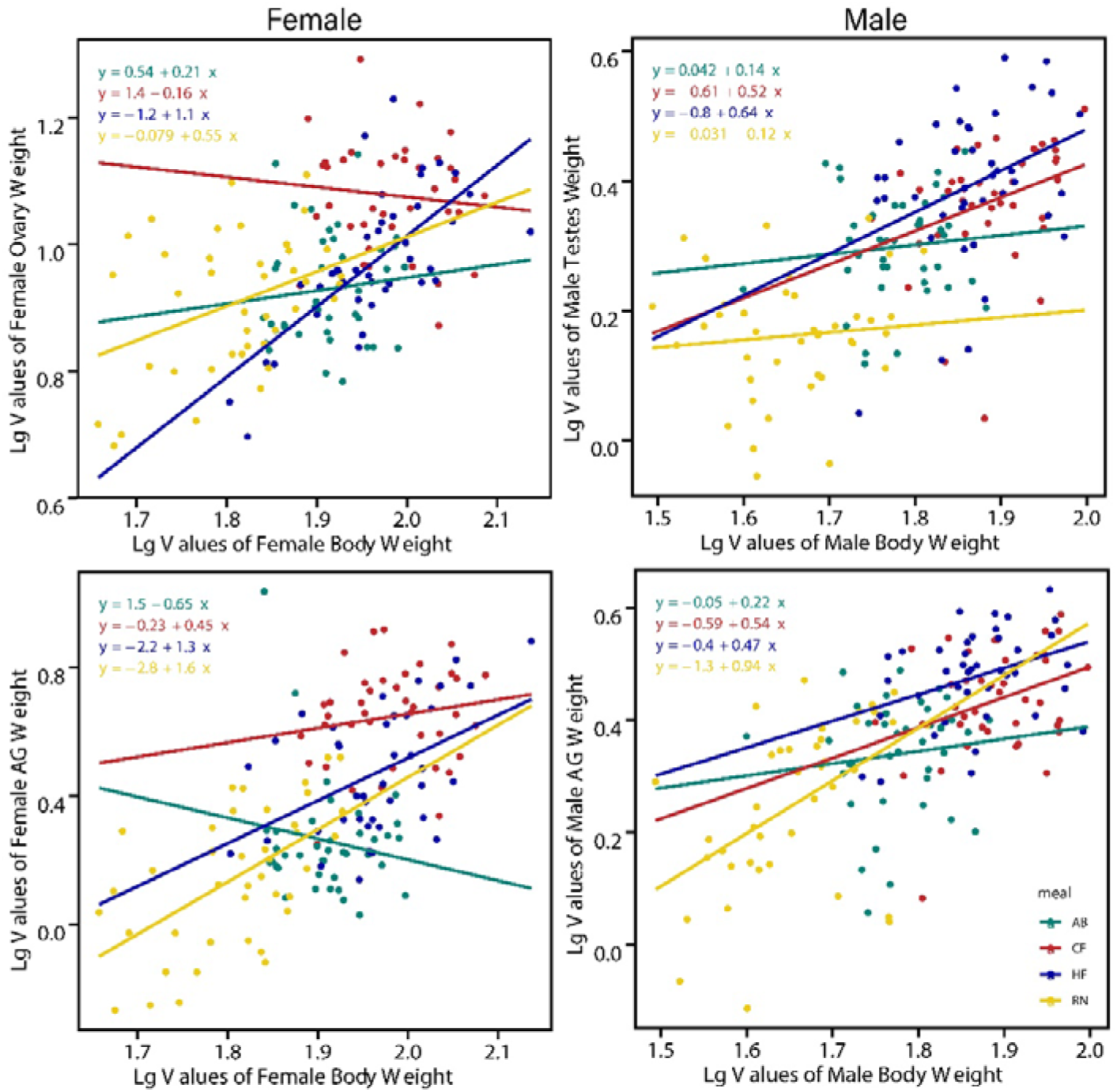
Allometric Scaling Relationships Between Body Weight and Reproductive Organ Weight in BSFs. The top panels display the correlation between body weight against ovary weight for females (left) and body weight against testes weight for males (right). The bottom panels show the relationship between body weight and accessory gland weight for females (left) and males (right). Each point represents an individual fly, with the different colours indicating the larval meal treatments. The fitted lines correspond to the linear regression for each treatment group, with the equations providing the allometric coefficient (slope) and intercept for each regression line.

### Diet-Induced Plasticity in Allometric Growth of Reproductive Organs

Figure 3 shows the allometric relationships of reproductive organs. In females, the allometric growth of ovaries varied across the different meal treatments. The allometric coefficient for RN, HF and AB were positive, while negative for CF (coefficients= -0.16) suggesting a reduction or a slower rate of increase in ovary weight relative to body weight when reared in CF. Only HF females exhibited a hyperallometric pattern with a coefficient of 1.1, suggesting a disproportional increase in ovary weight per unit increase in adult body weight when reared in HF. The allometric scaling of accessory glands to body size also exhibited plasticity. AB group showed a negative allometric coefficient (−0.65) in accessory gland development. A strong hyperallometric growth was observed in the RN treatment, with an allometric coefficient of 1.6, followed by HF at 1.3, both indicating that a disproportional increase in accessory glands with every unit increase in body weight. The CF treatment had a positive but lower coefficient of 0.45, indicative of a hypoallometric growth pattern. (Figure 3)

In males, both testes and accessory glands relative to body weight displayed hypoallometry (allometric coefficient < 1) across all dietary treatments. RN males exhibited the highest allometric coefficient for accessory glands (coefficients= 0.94), with CF and HF showing intermediate values (coefficients= 0.54 and 0.47), and AB the lowest (coefficients= 0.22). Conversely, testes size in RN males had the smallest allometric coefficient (coefficients= 0.12), the AB group was marginally higher (coefficients= 0.14), while males from the HF and CF diets had notably larger coefficients (coefficients= 0.64 and 0.52) (Figure 3).

### Diet-Induced Plasticity in Mating behaviours and Female Egg Production

Figure 4 illustrates that CF adults exhibited the longest mating latency across the first five mating events, with HF adults displaying the second-longest latency starting from the third event. In contrast, RN adults demonstrated the shortest mating latency. Males from the RN and HF dietary groups initiated the most mating attempts, with AB males following. CF males, however, exhibited notably fewer attempts, with their frequency being half that observed in males from the other groups. CF adults exhibited the least successful mating events, whereas RN adults achieved the greatest number, followed by AB. In contrast, HF adults, despite having the highest frequency of male mating attempts, displayed the lowest success in actual mating events. Asu such, HF had the lowest proportion of successful mating attempts, while RN males had the highest success ratio in mating attempts, succeeded by AB and CF (Table 3). No significant differences were observed in female resistance duration across the various groups. Likewise, the egg production per individual female did not vary significantly among the groups, although a non-significant trend was apparent with CF, HF, AB, and RN descending in the rank of the egg production weight.

**Figure 4.**
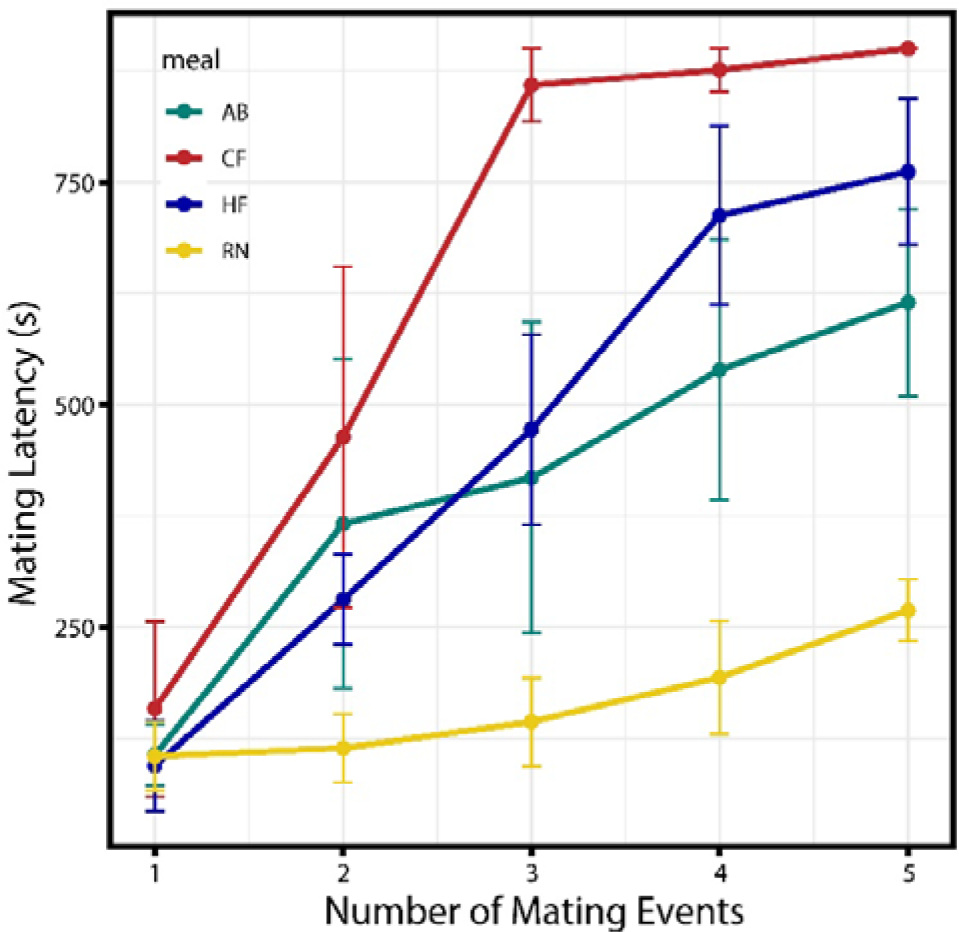
Mating latency of the Initial Five Mating Events. The latency was measured in seconds, defined as the duration from the start of the mating experiment to the establishment of a mating pair, for each discrete event. Different meals are categorized by colours. Error bars represent the standard errors from the four experimental replicates.

**Table 3.** Parameters of Mating Behaviours and Individual Egg Production in Females.

| Meal | Mating attempts | Successful mating events | Successful mating attempt ratio (%) | Female resistance duration (s) | Female egg production (mg) |
| --- | --- | --- | --- | --- | --- |
| AB | 65.45 ± 6.55 b | 6.30 ± 1.78 b | 9.73 ± 2.77 b | 5.75 ± 3.82 a | 21.64 ± 3.14 a |
| CF | 33.75 ± 10.07 c | 3.00 ± 1.07 d | 9.35 ± 2.90 bc | 6.50 ± 5.07 a | 27.90 ± 13.92 a |
| HF | 77.94 ± 6.92 a | 4.78 ± 1.11 c | 6.08 ± 1.15 c | 7.11 ± 8.81 a | 26.52 ± 1.44 a |
| RN | 70.84 ± 15.47 ab | 10.91 ± 1.29 a | 15.95 ± 2.96 a | 3.72 ± 1.93 a | 16.65 ± 3.97 a |

**Note:** The values represent mean ± standard error, and the letters behind the values represent significant difference according to TukeyHSD test and Kruskal test.

### Diet-Induced Plasticity in Longevity and Protandry Degree in BSF Adults

The survival analyses of BSFs as depicted in Figure 5a reveal significant differences in longevity across the dietary treatments for both sexes (Log-Rank Test, p < 0.0001). Overall, males displayed greater longevity than females, with males also showing increased plasticity in lifespan. Larvae reared on the RN diet tended to achieve the longest adult lifespan for both sexes, followed by those reared on HF, CF, and AB, in that order. Notably, the AB diet was associated with the shortest longevity among both female and male BSFs. The significance of protandry was affected by rearing meals (Figure 5b). In the early phase of adult emergence, diets RN, HF, and CF exhibited a male-dominant sex ratio pattern, suggesting precocious emergence of males relative to females in these groups. The RN diet was the most significant in this pattern, with the sex ratio equalized post day 20. The HF diet achieved a balanced sex ratio by day 10, while the CF diet reached within the first 5 days. In contrast, the AB diet did not exhibit this male-biased emergence pattern. (Figure 5b)

**Figure 5:**
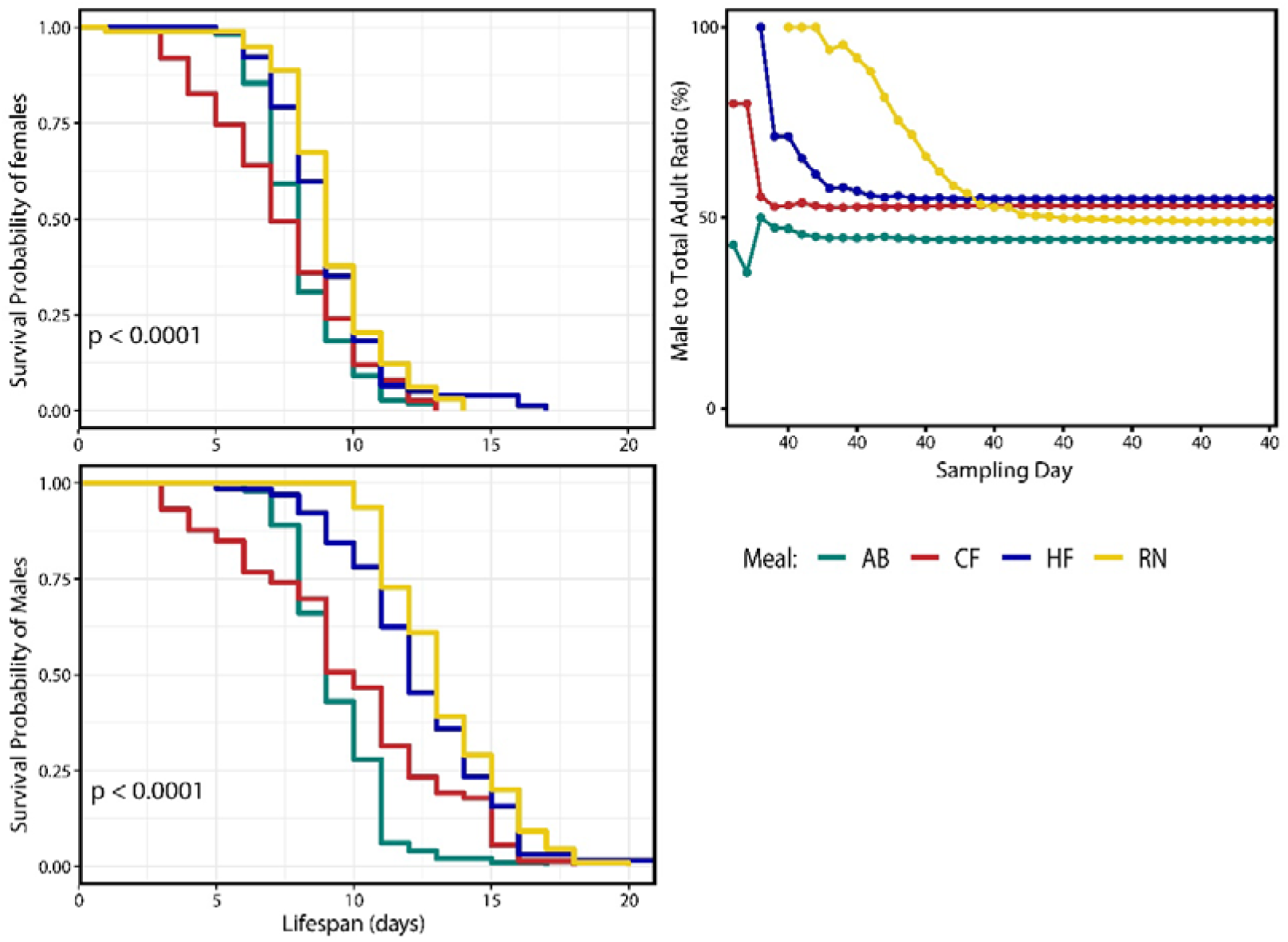
Influence of Larval Diets on Longevity and Protandry in Black Soldier Fly Adults. This figure displays: (a) Cumulative Survival Curves for both male and female BSFs under various dietary treatments over time (Log-Rank Test, p < 0.0001), showing the survival proportion at each sampling interval with different diets represented by distinct colours; and (b) Changes in the sex ratio of newly emerged adults (depicted as the percentage of males to total adults) over time under different larval dietary treatments (categorized by colours).

## Discussion

### Sexual Dimorphism in Phenotypic Plasticity

Previous studies have showed that when diet quality is manipulated, body size plasticity is greater in females than in males (Stillwell et al. 2010). In this study, males showed a slightly higher plasticity in body size, while for reproductive organ parameters (GSI and accessory gland to gonad ratio), females showed significantly higher degree of plasticity than males. The “plasticity-first evolution” theory suggests that plasticity can facilitate adaptive phenotypic change for selection to act on, while under long-term strong selection, canalization may be a more effective strategy as opposed to maintaining plasticity (Ghosh et al. 2019, Levis and Pfennig 2019). Males often experience stronger sexual selection on reproductive traits (Glaudas et al. 2020, Davies et al. 2023), thus the intense selection on BSF males may result in reduced plasticity in traits such as the accessory glands and testes in response to environmental variation. Overall, we note that females produce gonads that are much heavier than their accessory glands, but this pattern is reversed in males. Across all diets, the weight of male accessory glands was heavier than that of the testes.

Notably, male the accessory glands to testes ratio exhibited no plastic response with dietary treatments. This robustness to dietary variation implies that the relative investment in accessory glands and testes may tightly linked to male reproductive fitness and under strong selective pressure for an optimal ratio. Previous studies indicate that, in insects, the size of the male accessory glands can play a more crucial role in reproductive success than the size of the testes (Bangham et al. 2002, Baker et al. 2003). In males, these glands are responsible the production of seminal proteins that are often involved in facilitating sperm maturation and transfer to females (Avila et al. 2011). Additionally, the seminal proteins may serve as nutrient resources for females (Raikhel et al. 2005) or induce behavioural changes to mitigate sperm competition (McNamara et al. 2009). Larger testes suggest increase sperm production (Kahrl et al. 2019), which in turn could require a proportional increase in accessory glands to facilitate sperm transfer, fertilization, and egg development (Wigby et al. 2009). Therefore, maintaining an optimal accessory gland-to-testes size ratio may significantly impact the reproductive fitness of BSF males, evolving to resist environmental variations in this species.

### Diet-induced Morphological Plasticity in Body Size and Reproductive Organ Weight

Diet-induced morphological plasticity in body and reproductive organs has been demonstrated in many insect species (Nattero et al. 2013, Thompson 2019). Yet, few studies have focused on the ontogenetic plasticity of reproductive organs of BSF. Our study firstly provides data based on dissection to investigate diet-induced plasticity in BSF reproductive organ weight for both male and female. In our study, we observed distinct patterns of diet-induced morphological plasticity in body weight and reproductive organ weight among BSF adults. The trend followed the order CF ≥ HF > AB > RN, indicating that diet type significantly influences these morphological traits. CF, often used as a control substrate in numerous BSF studies, is recognized as an ideal diet for BSF larvae (Bosch et al. 2020), leading to larger larvae with more substantial nutrients (Lalander et al. 2019, Zhang et al. 2023b). Interestingly, our study suggests that the long-term meal adaptation of BSFs to the HF diet since 2018 may have played a crucial role in enhancing larval growth, consequently resulting in larger adult sizes and heavier reproductive organs.

Notably, HF males exhibited the heaviest reproductive organ (comparable to that of CF males), despite HF males having shorter scutum lengths than CF males. This result may indicate that meal adaptation can have different impacts on males and females; particularly, it appears to have more pronounced effects on the reproductive organ development in males and on body size development in females. This finding aligns with previous research that highlights adaptation can be sexually dimorphic (Connallon 2015). Specifically, different sexes may reach distinct fitness optima for shared traits during meal adaptation, potentially leading to sexually antagonistic genetic variance (Delcourt et al. 2009). The sex-specific differences in response to long-term meal adaptation observed in our study may reflect such an evolutionary process that involves optimizing distinct traits between the sexes to enhance fitness. Further research is needed to understand the underlying genetic and physiological mechanisms of this sexually dimorphic adaptation.

### Diet-induced Morphological Plasticity in Sperm Length and Sperm Allometry

Previous studies, limited by smaller sample sizes, have reported significant differences in BSF sperm lengths. For instance, Munsch-Masset et al. (2023) analysed six sperms from four males and reported a total sperm length exceeding 3600 μm, with sperm head length averaging 16.2 ± 1.04 μm. In an analysis of six individual males, total sperm lengths around 860 µm and sperm head lengths near 8 µm were observed (Malawey et al. 2019). Additionally, Kotzé et al. (2019) noted sperm head lengths ranging from 9 to 21μm in a single male. In contrast, our larger sample size yields more comprehensive data, showing that total sperm lengths range from approximately 500 to over 1000 μm, and sperm head lengths from about 15 to 21 μm within the same dietary treatment group (Figure 3c). Currently, there is no evidence to suggest that BSF exhibits sperm polymorphism, a trait often seen in other insect species. Therefore, the adaptive significance of this observed variation in sperm length in BSF is a compelling subject for further investigation.

In contrast to many studies on insects that have focused on the effects of diet nutrition on sperm quantity (sperm number) (Abraham et al. 2011, Dávila and Aron 2017, Huck et al. 2021), this study investigates the influence of diet on sperm quality (sperm length). Early-stage environmental conditions for insects can impact sperm production (Liu et al. 2021), and our study suggest that larval diet can influence sperm length variation. The RN males, characterized by their smaller body size and reproductive organs, displayed shorter sperm lengths than males from other groups. This observation contrasts with prior research by Gage & Cook (1994), which indicated that a restrictive larval diet typically leads to a reduction in sperm number rather than a change in sperm size (Gage and Cook 1994). In our study, the egg production of mated females showed no significant differences across the various diet groups. In addition, a study on dung beetles indicated that fertilization success was biased toward males with relatively shorter sperm (García-González and Simmons 2007). Therefore, the observed reduction in sperm length among RN males might suggest an adaptive strategy aimed at enhancing reproductive success.

The CF males, characterized by their low body protein content (Zhang et al. 2023b), exhibited a significant positive allometric growth in sperm length relative to body size. This observation is aligned with findings from a previous study on ants, which indicated that protein restriction influences sperm count but not quality (Dávila and Aron 2017). Notably, in the RN group, we observed that smaller males tended to have longer sperm lengths, a trend that aligns with the negative association reported in a study on scorpion species (Vrech et al. 2014). Future research could focus on elucidating the trade-off between sperm length and number in BSF males exposed to varying environmental conditions.

### Impact of Diet on Mating Behaviours and Egg Production in BSF

RN adults, despite being the smallest in body size and reproductive organ size, demonstrated the best mating behaviour performance: the shortest mating latency, the highest success rate in mating attempts, and the most numerous total mating events. This contrasts with CF and HF adults, though larger in size and with heavier reproductive organs, exhibited longer mating latencies and fewer successful mating events.

In the insect world, the cost of accurately identifying the sex of potential mates, or the associated cost of hesitation, often exceeds the cost of making erroneous mating attempts (Scharf and Martin 2013). Thus, male-male sexual interactions have been observed in many species including BSF (Scharf and Martin 2013, Giunti et al. 2018). In our study, we observed that male BSFs engaged in mating attempts with both males and females, resulted in successful mating attempt ratio less than 100%. In addition, plasticity in mating attempt accuracy was observed in results, with the smallest RN males showing the highest ratio of successful mating attempts. The smaller RN adults possess relatively fewer energy resources, leading to a higher energy expenditure per mating attempt (Pitnick and Markow 1994, Hayward and Gillooly 2011). Consequently, precision in mating attempts becomes crucial for them, leading to a significantly higher ratio of successful mating attempts.

The mating behaviours of adult BSFs has been largely understudied, despite its significance in revealing reproductive mechanisms for improved BSF cultivation (Julita et al. 2020, Meneguz et al. 2022, Lemke et al. 2023). Our research is the first to demonstrate that larval diet can induce plastic responses in mating latency and mating attempt accuracy. Importantly, our findings reveal that larvae reared on a less suitable diet (RN), which results in smaller larvae and adults, appear to compensate for this growth disadvantage by optimizing their mating behaviours. This optimization, characterized by reduced mating latency and greater mating attempt accuracy, might enable RN adults to achieve reproductive fitness comparable to individuals from other dietary groups.

The absence of variation in female resistance duration across dietary treatments suggest that female receptivity may be less affected by diet than male mating behaviours. Moreover, the weight of individual female egg production showed no significant variation, aligning with a previous study where rearing wild-caught BSFs on different diets did not result in plasticity in individual female egg cluster sizes (Tomberlin et al. 2002). Given the critical role of female fecundity in the overall reproductive fitness of the population (Bradshaw and McMahon 2008), the stability of female egg production could indicate an adaptiveness to maintain reproductive success.

Overall, these findings underscore the intricate connection between diet, mating behaviour, and potential reproductive outcomes in BSFs. Future studies on the physiological and molecular mechanisms driving these diet-induced changes in mating behaviours could provide valuable insights into how BSFs adapt to different nutritional environments, a line of inquiry that holds significant implications for optimizing BSF cultivation and management.

### Plasticity in Allometric Growth of Reproductive Organs

For BSF adults, each individual has a given amount of resource stored from the larval stage, and it allocates these resources to different structures. Trade-offs involved in resource allocation often result in the prioritization of certain structures, like reproductive organs, potentially at the expense of others (Weiner 2004). Allometry, in this context, quantitatively describes the relationship between growth and the distribution of resources (Cheplick 2005). In our study, the allometric growth patterns observed in BSFs reveal distinct differences between the sexes, which may indicate sex-specific adaptive strategies. In females, the allometric growth of ovaries and accessory glands showed more significant variability across different larval diets, indicating a higher degree of phenotypic plasticity. This variability indicates that female reproductive organ development is particularly sensitive to nutritional variations experienced during the larval stages. Conversely, male BSFs exhibited a more consistent hypoallometric growth pattern for both testes and accessory glands, regardless of the larval diet. Considering that BSF adults do not require feeding to perform all mating activities (Gobbi et al. 2013), their energy resources are inherently limited. Consequently, an optimized strategy for energy allocation becomes crucial. This optimization is essential not only for facilitating active mating activities (function of non-reproductive body parts), but also for ensuring effective fertilization (function of the reproductive organs) maximize total reproductive fitness. Additionally, males have historically been subject to stronger sexual selection forces compared with females (Glaudas et al. 2020, Davies et al. 2023). This intense sexual selection pressure could play a significant role in shaping their energy allocation strategies and lead to a reduced plasticity in allometric growth pattern for their reproductive organs compared to females.

Females are often more sensitive to nutritional stress during the larval stage than males (Teder and Kaasik 2023). This is reflected in our study, where the allometric scaling of female reproductive organs showed more significant plasticity, suggesting that the female reproductive organ development is particularly sensitive to nutritional variations experienced during the larval stages. CF females showed inverse allometric growth (allometric coefficient < 0) for ovary growth. Prior research indicates that larvae reared on the CF diet have a significantly low protein content (Zhang et al. 2023b), a vital component for insect ovarian development (Wei et al. 2017, Sun et al. 2022). This protein deficiency in CF females may constrain the proportional growth of ovary weight with body weight for females, leading to the inverse allometry observed in our study. An alternative explanation could relate to developmental constraints of the organ (Voje et al. 2014). HF females, despite similar body sizes to CF females, had smaller ovaries and ovary weight increased faster than body weight. This may suggest that in CF females, ovary size might have reached a developmental limit, contributing to the inverse allometric pattern observed.

The plasticity in the allometric growth of accessory glands in females further highlights the influence of diet on reproductive organ development. RN and AB females exhibited comparably lower body and accessory gland weights than CF and HF females. However, the allometric growth pattern of accessory glands in RN and AB females were distinctly different. RN females exhibited a robust hyperallometric growth for accessory glands, while AB females showed an inverse allometric relationship in their accessory gland growth (allometric slope<0). In insects, fluid from female accessory glands is vital for nourishing sperm, aiding fertilization, and providing protection during mating, thereby boosting reproductive success (Avila et al. 2011). Within a population, fecundity typically correlates with female body size; larger females tend to attract more males and engaged in more mating activities to acquire enough sperm for egg fertilization (Kokko and Jennions 2023). This trend is especially pronounced in polygynandrous species like BSFs. Our study observed that RN females engaged in mating activities more frequently than AB females, suggesting that larger RN females might need proportionally larger accessory glands to effectively facilitate fertilization and provide protection during mating events.

In males, the growth of both testes and accessory glands relative to body weight displayed hypoallometry (allometric coefficient < 1) across all dietary treatments. This pattern, where male reproductive organ growth is proportionally less than body growth, aligns with the “one size fits all” hypothesis (Eberhard et al. 1998), suggesting that male genitalia adapted to match the most common female size enhances mating success. As non-feeding adult flies, such hypoallometric growth in males may optimize reproductive fitness by ensuring compatibility with female organs, while also conserving energy by not over-investing in reproductive organ development.

### Correlated Plasticity in Protandry and Longevity as a Reproductive Strategy

Protandry, a phenomenon commonly observed in various insect species, including BSF (Tomberlin et al. 2002, Harjoko et al. 2023), has traditionally been considered a stable population trait unaffected by environmental changes (Fischer and Fiedler 2000, Zijlstra et al. 2002). However, our study reveals the degree of protandry can exhibit plasticity. Notably, observed that males exhibited greater plasticity in longevity compared to females, and individuals exhibiting extended longevity also displayed increased protandry degree. This may suggest a potential combined strategy in the reproductive dynamics of BSF population when facing dietary environmental changes. In BSF populations, males generally have a longer lifespan than females (Harjoko et al. 2023). Therefore, in populations with extended longevity, it is advantageous for males to emerge before females, as this early emergence strategy increases their opportunities for mating throughout their lifespan. Conversely, in populations where longevity is reduced, if males continue to emerge significantly earlier than females, it can result in under-utilization of mating resources. This is due to males with shorter lifespan, emerging earlier, tend to die before females, resulting in a limited lifespan overlap between the sexes.

Consequently, in our study, a trend was observed where groups with shorter lifespans displayed reduced protandry degree, while those with extended lifespans exhibited more significant protandry. This adaptive strategy could maximize the lifespan overlap between males and females, thereby optimizing mating success in various environmental conditions. It was reviewed that within protandrous species, increased protandry degree is associated with increased female development time (Teder et al. 2021), and a mathematical model predicts that protandry and reproductive lifespans increase as environmental stress decreases (Morbey and Abrams 2004). However, our study appears to be the first to formulate a hypothesis regarding the interaction between male longevity and protandry degree and its potential impacts. Based on our findings, we hypothesize that in populations where males have a longer lifespan than females, the relationship between male longevity and protandry degree significantly influences the lifespan overlap between sexes. Specifically, our hypothesis suggests that in cases of shorter male lifespans, reducing the degree of protandry is an adaptive response to enhance their lifespan overlap with females. (Figure S4 elaborates on this protandry-lifespan hypothesis)

## Conclusion

This comprehensive study on Black Soldier Fly (BSF) adults has revealed key insights into the phenotypic plasticity of reproductive traits influenced by dietary variations. First, we observed a markedly reduced plasticity in reproductive organ parameters in BSF males, due to stronger sexual selection pressures. Importantly, a consistent accessory gland to testes ratio was noted across all diets, emphasizing its significance in male reproductive fitness. Secondly, our study highlighted the plasticity in protandry and longevity as a potential reproductive strategy. The observed correlation between increased male longevity and protandry degree suggests an adaptive strategy to optimize lifespan overlap between the sexes, thereby enhancing mating success under various environmental conditions. Moreover, our results show that larval diet has a lasting impact on adult size and reproductive organ development. Meal adaptation appears to affect males and females differently, with more pronounced effects on reproductive organ development in males and body size in females. Additionally, our study provides fresh insights into how diet influences sperm length, highlighting the potential adaptive significance of variations in sperm length and allometry. Our research also indicates that diet can induce plasticity in adult mating behaviours: larvae reared on less suitable diets, resulting in smaller adults, seem to compensate for this disadvantage by optimizing mating behaviours to enhance reproductive success. Notably, BSF females displayed greater plasticity in the allometric growth of reproductive organs, while BSF males consistently exhibited negative allometric growth, aligning with the ’one-size-fits-all’ hypothesis.

Overall, our research underscores the complex interplay between diet, reproductive traits, life history traits, and mating behaviours in BSFs, revealing the significant role of dietary conditions in shaping reproductive strategies. These findings pave the way for future research, particularly in unravelling the genetic and developmental pathways that drive the observed variations in allometric relationships and reproductive adaptations in BSFs.

## Acknowledgements

We would like to express our sincere thanks to the members of the Food Science and Technology Lab – Dr. Mei Hui Liu, Chin Wee Heng, and Koh Rui En – for conducting the nutrient analysis on the larval meals used in this study. Additionally, we are grateful for the insightful contributions of Nicole Lee, Tyrone Tan, Shaktheeshwari Silvaraju and fellow Reproductive Evolution Lab members that have enhanced this manuscript. This research was jointly funded by the National Research Foundation of Singapore (NRF2020-THE003-0003/A00085140000) as well as the Ministry of Education Singapore (A00044240000).

## Data availability

The datasets, videos and R codes generated and analysed during this study are available in the [BSF_Adult_Plasticity] repository, https://github.com/ReproLab/BSF_Adult_Plasticity.

## Supplementary Materials

**Figure S1.**
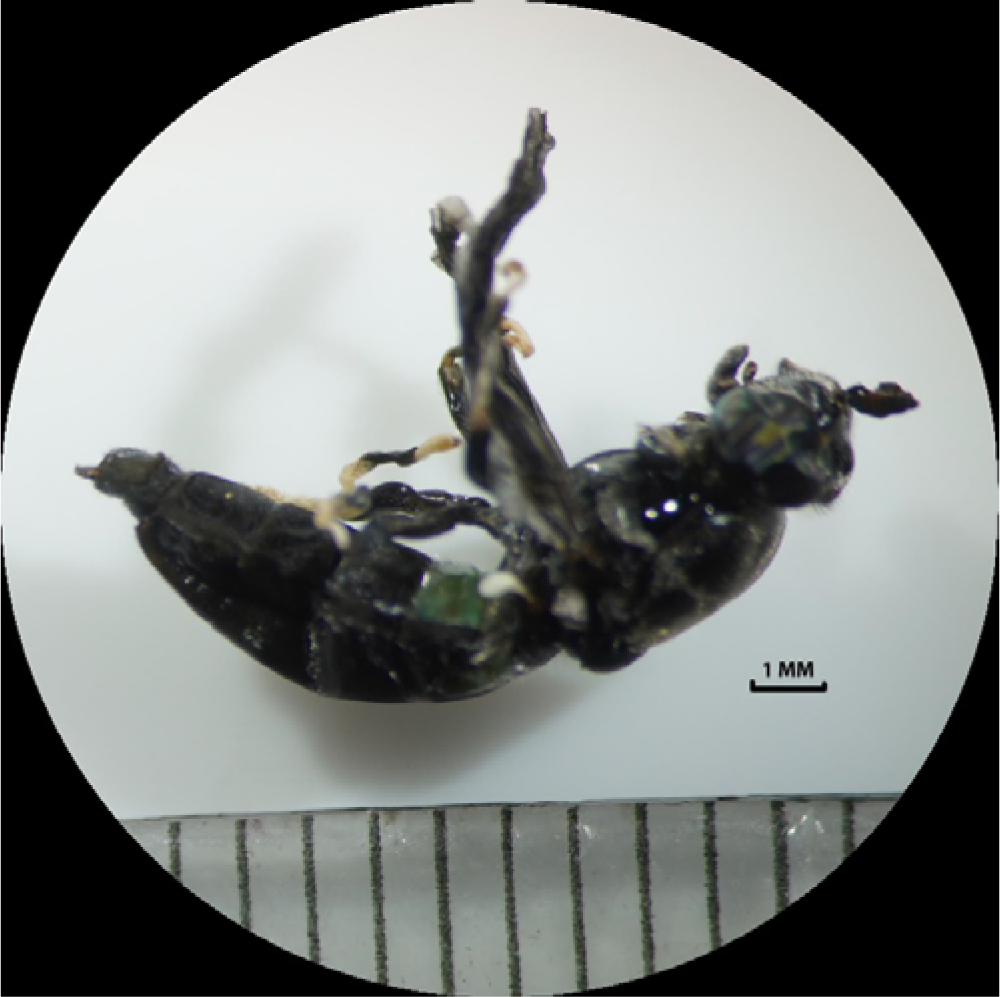
Abnormal Adult Individual From Larvae Reared on VF meal. As we observe, the abnormal including, incomplete wings, incomplete emerge, smaller size, twisted body shapes, and soft body.

**Figure S2.**
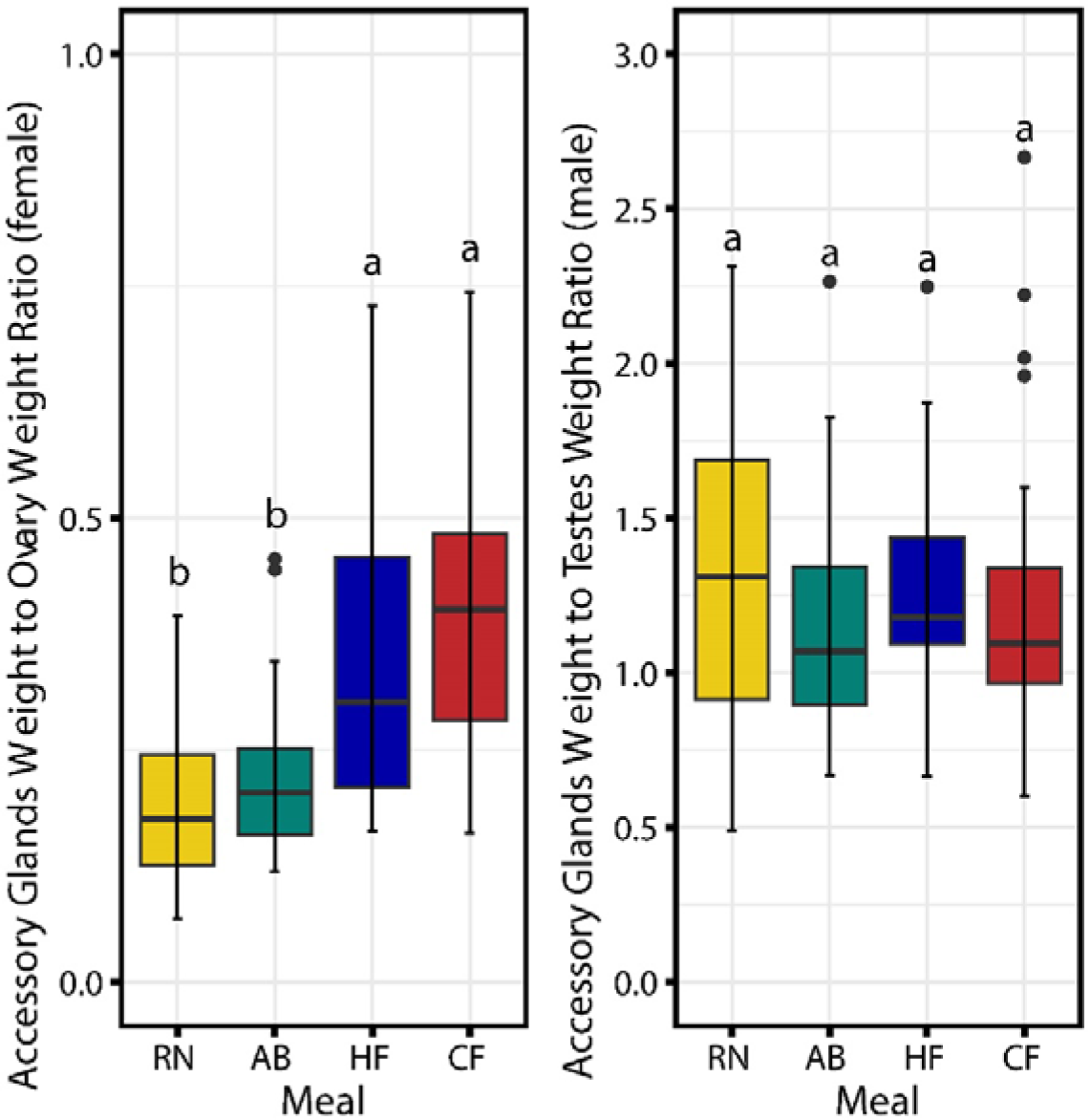
Accessary glands to Gonads (ovaries/ testes) ratio. (a) Ratio of Accessory Glands Weight to Ovary Weight; (2) Ratio of Accessory Glands Weight to teste Weight. Different meals are categorized by colours. Significant differences in ratios among meal treatments are indicated by lowercase letters (Kruskal-Wallis test, Bonferroni correction, P < 0.05).

**Figure S3.**
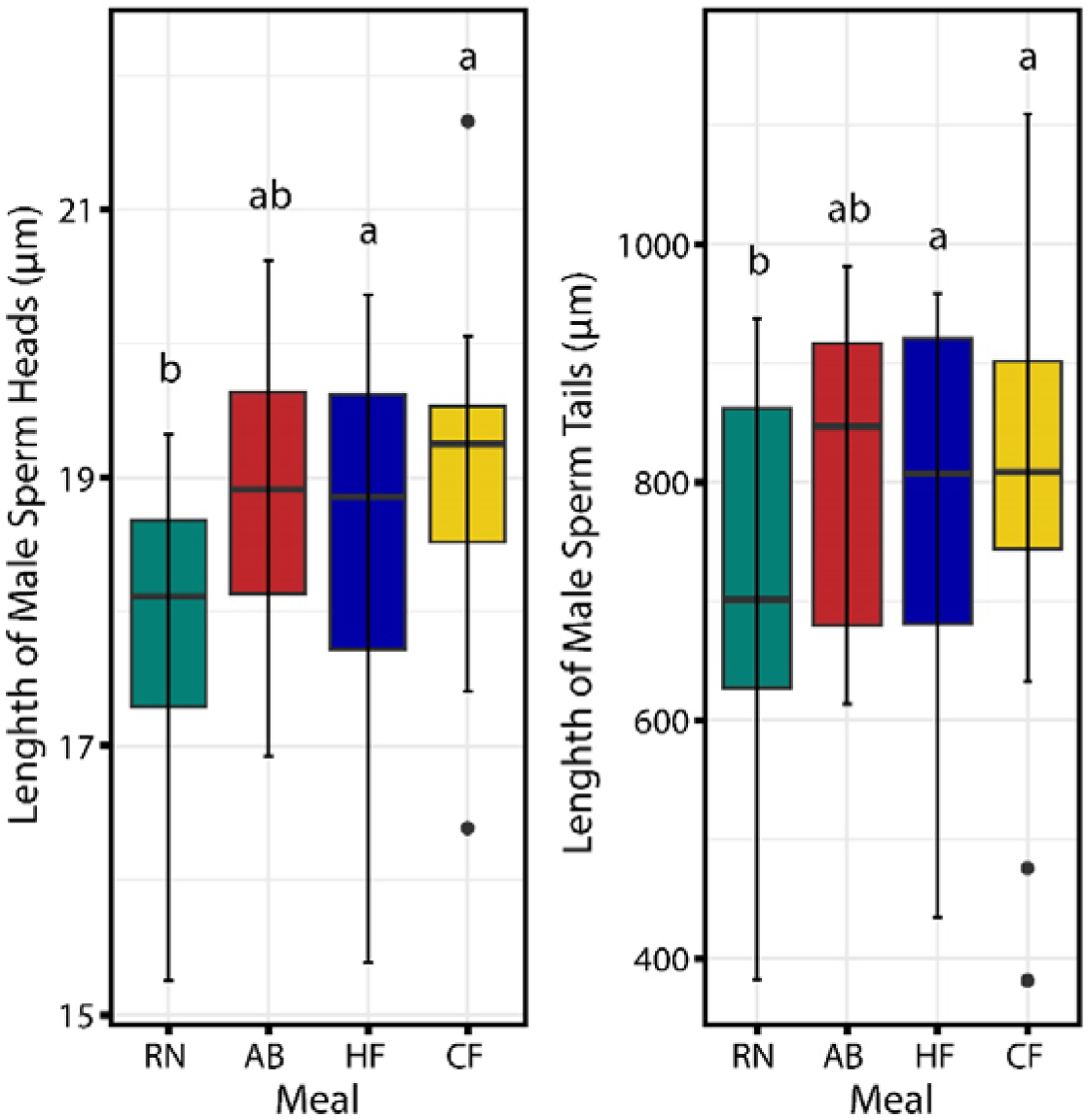
The length of Sperm Heads and Tails in BSF Males. Different meals are categorized by colours. Significant differences in lengths among meal treatments are indicated by lowercase letters (Kruskal-Wallis test, Bonferroni correction, P < 0.05).

**Figure S4.**
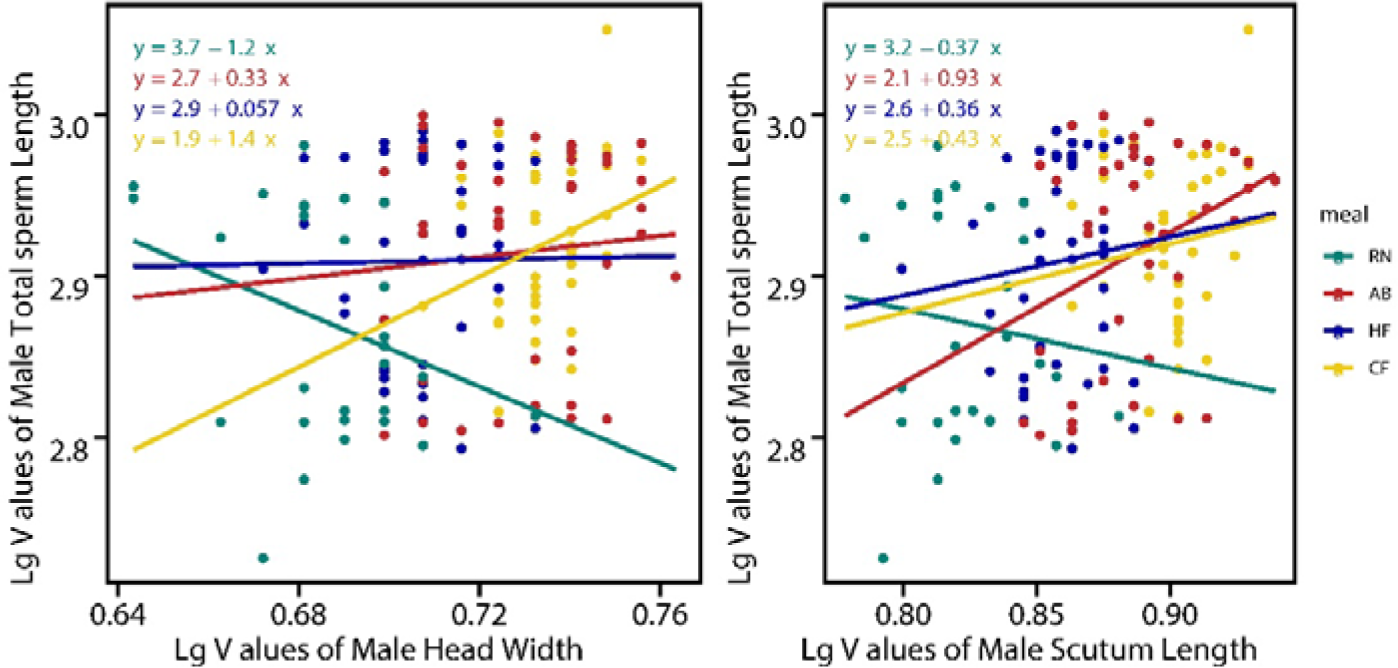
Allometric Scaling Relationships Between Total Sperm length and Body Size in BSF Males. (a) The correlation between total sperm length against head width for males; (b) The correlation between total sperm length against scutum length for males. Each point represents an individual fly, with the different colours indicating the larval meal treatments. The fitted lines correspond to the linear regression for each treatment group, with the equations providing the allometric coefficient (slope) and intercept for each regression line.

**Figure S5.**
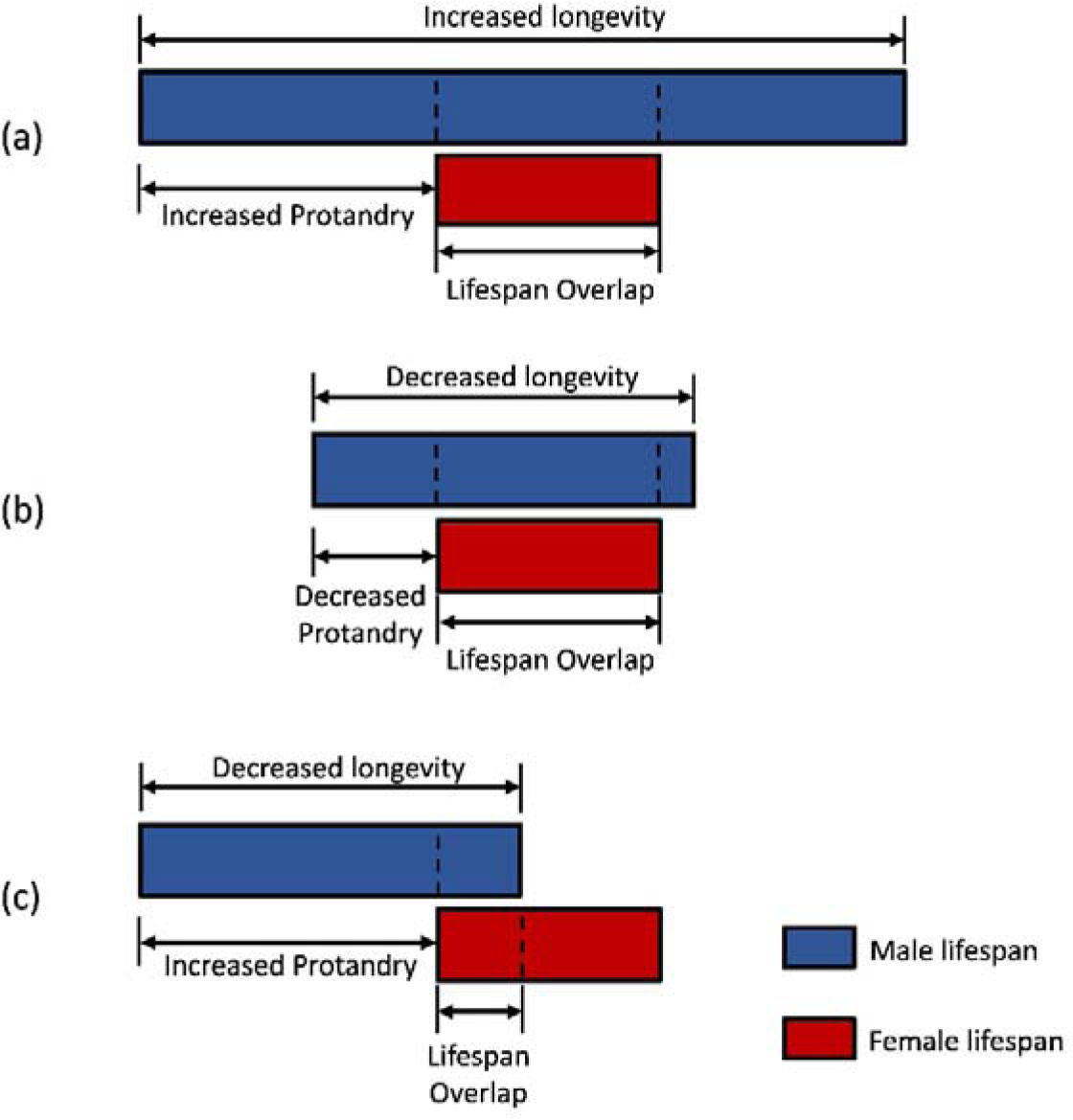
The Protandry-Lifespan Overlap Hypothesis Derived from This Study. This hypothesis posits that in populations where males outlive females, the interaction between longevity and protandry degree impacts the lifespan overlap between males and females. (a) Males with increased longevity and increased protandry degree resulted in maximized lifespan overlap; (b) Males with decreased longevity and decreased protandry degree resulted in maximized lifespan overlap; (c) Males with decreased longevity and increased protandry degree resulted in reduced lifespan overlap.

**Table S1-1.**
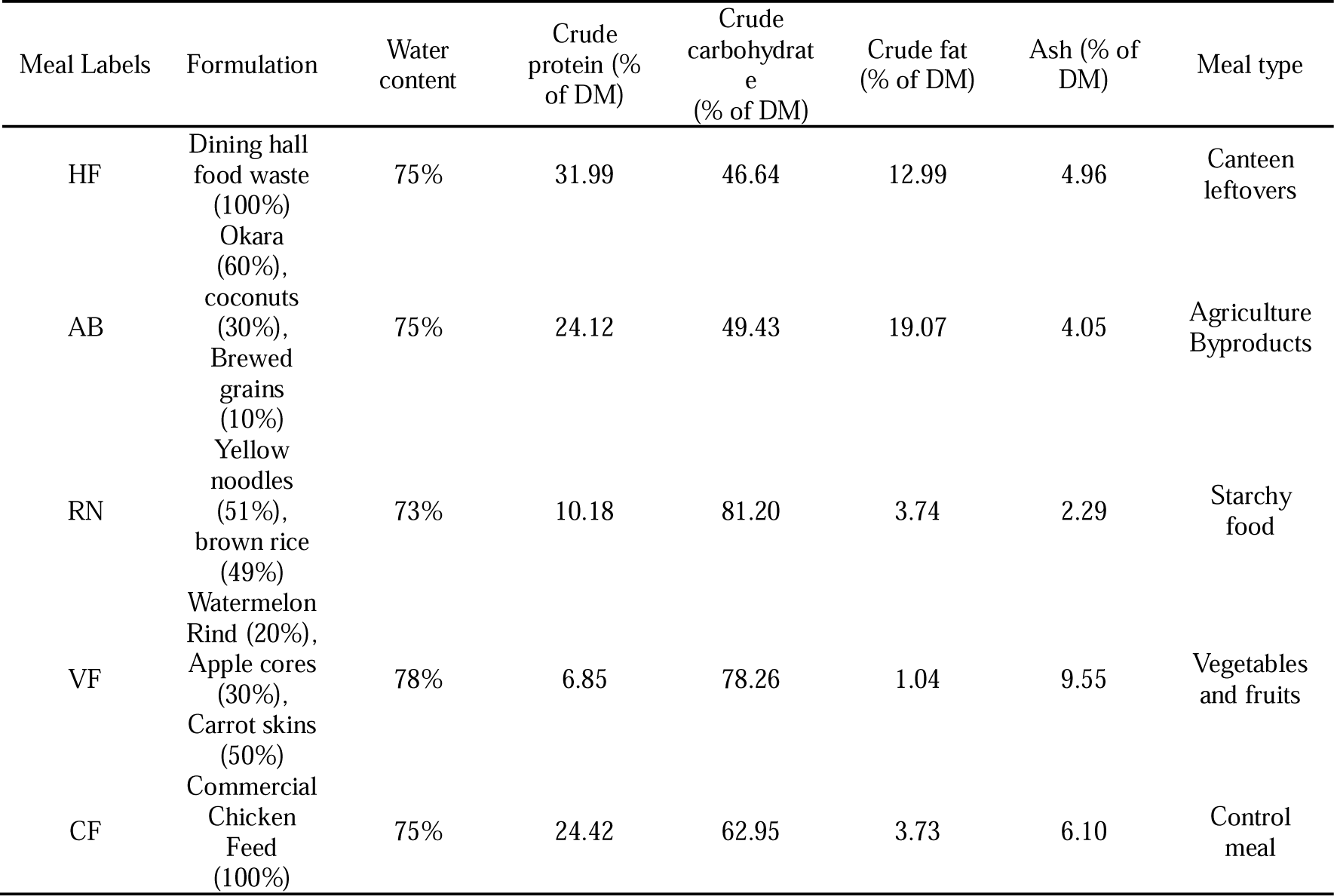
Macronutrient composition, formulation, and meal type of different larval meals.

**Table S1-2.** Amino acid profile of different larval meals.

| Amino Acid | HF (mg/100g) | AB (mg/100g) | RN (mg/100g) | VF (mg/100g) | CF (mg/100g) |
| --- | --- | --- | --- | --- | --- |
| Asp | 102.95 | 98.20 | 47.49 | 18.40 | 175.95 |
| Thr | 36.73 | 38.48 | 15.99 | 8.37 | 71.76 |
| Ser | 88.90 | 47.36 | 27.08 | 10.19 | 91.74 |
| Glu | 147.97 | 166.60 | 108.74 | 46.64 | 302.48 |
| Gly | 57.73 | 56.50 | 25.69 | 8.47 | 78.03 |
| Ala | 60.94 | 66.54 | 32.03 | 8.95 | 81.73 |
| Cystine | 47.51 | 15.24 | 16.02 | 1.81 | 15.64 |
| Val | 79.20 | 73.52 | 33.51 | 9.76 | 74.30 |
| Met | 32.96 | 43.86 | 54.33 | 7.09 | 42.02 |
| Ile | 44.74 | 65.82 | 25.90 | 9.44 | 76.43 |
| Leu | 96.94 | 97.46 | 46.69 | 15.83 | 146.22 |
| Tyr | 55.93 | 41.01 | 20.68 | 8.61 | 73.13 |
| Phe | 54.33 | 58.37 | 25.68 | 9.91 | 94.42 |
| Lys | 95.88 | 59.00 | 23.58 | 13.07 | 123.08 |
| NH<sub>4</sub>Cl | 0.32 | 0.39 | 0.21 | 0.08 | 0.46 |
| His | 23.36 | 24.33 | 11.76 | 7.01 | 55.71 |
| Arg | 82.80 | 70.07 | 34.34 | 13.14 | 118.38 |
| Pro | 49.23 | 53.49 | 45.94 | 11.26 | 90.96 |

**Table S2.** Effects of different larval diet on larval development parameters

| Parameters | Larval Meal |  |  |  |  |
| --- | --- | --- | --- | --- | --- |
|  | AB | CF | HF | RN | VF |
| Prepupa weight (mg) | 190.46 ± 4.22 b | 240.33 ± 5.53 a | 227.62 ± 7.17 a | 135.30 ± 4.01 c | 92.04 ± 4.53 d |
| Day 12 Prepupa number | 5.00 ± 4.36 b | 143.75 ± 10.08 a | 0 b | 0 b | 0 b |
| Day 13 Prepupa number | 34.25 ± 18.63 b | 175.75 ± 12.09 a | 13.00 ± 1.08 b | 0 c | 0 c |
| Day 14 Prepupa number | 70.5 ± 28.17 b | 202.0 ± 8.61 a | 50.0 ± 8.90 bc | 0 c | 0 c |
| Day 15 Prepupa number | 135.75 ± 30.17 b | 238.75 ± 7.79 a | 142.25 ± 10.04 b | 0 c | 0 c |
| Day 16 Prepupa number | 180.25 ± 26.16 b | 253.75 ± 12.24 a | 205.25 ± 9.43 ab | 0 c | 0 c |
| Day 17 Prepupa number | 203.75 ± 25.05 ab | 256.00 ± 12.86 a | 248.00 ± 12.02 a | 0 c | 22.50 ± 4.50 bc |
| Day 18 Prepupa number | 219.75 ± 26.31 ab | 256.50 ± 13.28 a | 281.00 ± 11.02 a | 7.00 ± 1.08 c | 73.50 ± 8.29 bc |
| Day 19 Prepupa number | 230.25 ± 25.46 b | 256.75 ± 13.49 ab | 303.25 ± 9.94 a | 20.00 ± 4.65 d | 155.75 ± 15.17 c |

**Note:** The values represent mean ± standard error, and the letters behind the values represent significant difference according to TukeyHSD test and Kruskal test.

